# Excitability and synaptic transmission after vitrification of mouse corticohippocampal slices

**DOI:** 10.1101/2024.06.03.597218

**Authors:** Alexander German, Enes Yağız Akdaş

## Abstract

Cryopreservation of adult neural tissue is of considerable practical and theoretical interest. Utilizing 61% w/v ethylene glycol, we vitrified and rewarmed acute mouse corticohippocampal slices to evaluate field excitatory postsynaptic potentials (fEPSP) in the stratum radiatum of the CA1 region of the hippocampus. Our results demonstrate successfully recovered synaptic transmission, and high-frequency stimulation (HFS)-induced potentiation. However, we failed to induce a stable potentiation following HFS stimulation. Structural analysis post-vitrification revealed cellular alterations such as swelling and vacuolization, which likely contributed to the unstable potentiation. Despite high variability in results, this study highlights the potential of vitrification to partially preserve brain function.

## Introduction

Luyet and Gehenio initially motivated cryobiology as a tool to investigate the nature of life [1]. Indeed, cryoimmobilization in the vitreous state later became a cornerstone of structural biology [2, 3]. Reversibly arresting brain tissue in the vitreous state might serve theoretical inquiries regarding the nature of mind [4], in addition to practical applications in neuroscience.

Since the discovery of the neuron [5, 6], microscopic investigations into the structure of brain tissue have continually gained in scope and resolution [7, 8], culminating in synaptic-resolution connectomics [9, 10]. The effort of mapping a mouse connectome [11] might contribute to neuroscientific understanding [12], especially if the structural data permits prediction of functional behavior. Acquiring such structural data presently relies on the ability to immobilize brain tissue in its native state. While small specimens like roundworms can be arrested in the vitreous state [13, 14], larger specimens like the fruit fly and mouse brain have been fixated with aldehydes instead [15, 16]. Deviations from the native organization after aldehyde fixation are manyfold [17], including a marked loss of extracellular space [18, 19]. Therefore, reversible vitrification of brain tissue could be an alternative to get closer to the native state of brain tissue.

In 1953, Luyet and Gonzales demonstrated the possibility to cryopreserve embryonic avian brain tissue using 60% ethylene glycol [20], and this possibility was later shown for fetal rodent and human brain tissue as well [21-24]. To our knowledge, full electrophysiological recovery after cryopreservation of adult mammalian brain tissue or neurons has thus far not been demonstrated [25, 26]. Notably, some recovery of activity was reported in the adult feline brain after several years at -20°C storage [27, 28], and in rodent ganglia after up to 24 hours at -76°C storage [29], both using 15% glycerol.

Pichugin et al. have demonstrated recovery of K(+)/Na(+) ratios after vitrification of rat hippocampal slices using 61% w/v VM3 [30]. Here, we vitrified mouse corticohippocampal slices using 61% w/v ethylene glycol (EG) and investigated the electrophysiological recovery of field excitatory postsynaptic potentials (fEPSP) in the stratum radiatum of CA1 subfield in response to stimulation of the Schaffer collaterals (SC).

## Materials and methods

This study used male and female C57BL-6N mice aged 10-13 weeks. The mice were maintained under standard laboratory conditions, which included a 12/12 light/dark cycle with lights on at 06:00, a temperature of 22°C ± 2/4, and 60% humidity, with water and food available ad libitum. All experiments were conducted in accordance with the guidelines for the Care and Use of Laboratory Animals set by the Government of Unterfranken and the European Communities Council Directive (2010/63/EU).

Animals were sacrificed by cervical dislocation and the brains were immediately transferred into ice-cold carbogenated (95% O_2_, 5% CO_2_, Carbogen Lab, Linde Gas, Germany) calcium-free artificial cerebrospinal fluid (aCSF, composition: 129 mM NaCl, 21 mM NaHCO3, 1.25 mM NaH2PO4, 1,8 mM MgSO4, 10 mM KCl, 10 mM glucose, 300+-10 mosmol). 400 μm-thick acute horizontal slices consisting of entorhinal cortex and hippocampus were prepared using Leica VT1200 Vibratome (Leica Biosystems, Germany).

The cortico-hippocampal region was dissected inside the aCSF and transferred onto a Netwell polyester mesh insert (Corning, USA) for 6-well plates. 6-well plates were positioned on the surface of a water bath inside a polystyrene box that was set to 10°C by mixing tap water and ice. Using the mesh insert, the corticohippocampal slices were immersed into 10°C solutions of increasing concentration of EG in a LM5 carrier solution. LM5 consists of 90mM glucose, 45mM mannitol, 45mM lactose, 8.2mM KCl, 7.2mM K2HPO4, 5mM reduced glutathione, 1mM adenine and 10mM NaHCO3. EG loading of slices was performed in 0%, 2%, 4%, 8%, 16% and 30% w/v EG for 10 minutes each. Subsequently, the mesh insert carrying the slices was transferred into three iterations of the full vitrification solution of 61% w/v EG for 10 minutes each. These solutions were set to -10°C in a second polystyrene box containing a 65% w/v EG in water bath that was kept at -10°C using dry ice. Subsequently, the mesh insert was transferred into liquid nitrogen, and slices were kept at -196°C for at least 10 minutes and up to 20 hours. Slices were rewarmed by rapidly transferring the mesh insert into 100 ml vitrification solution at -10°C. Subsequently, cryoprotectant unloading of slices was performed by increasing the osmolarity of the LM5 carrier solution by adding 300mM mannitol and immersion in 30% w/v EG at -10°C for 10 minutes, followed by immersion in 16%, 8%, 4%, 2%, 0% EG at 10°C for 10 minutes each, followed by a final step of LM5 without added mannitol. All chemicals were obtained from Sigma-Aldrich, USA.

The slices were extracted from the mesh insert and transferred to a Haas-type interface chamber with continuous aCSF perfusion and kept at 32°C for 2 h to recover prior to the recordings. Throughout recovery and recordings, slices were kept under continuous perfusion of carbogenated aCSF with a flow rate of 1.48-2 ml/min. Glass sharp electrodes (GB150EFT-10, Science products, Germany) that were used for recording, were prepared using a Flaming/Brown type micropipette puller (P-1000, Sutter Instruments, Germany). A stimulation electrode with 75μm tip separation and 0.1 MΩ impedance (WE3ST30.1B3, MicroProbes for Life Science, USA) was positioned at the SC, ensuring that the collaterals remained in the middle of the two tungsten tips of the electrode. The recording electrode was filled with aCSF (tip resistance 2-3 MΩ) and placed in the stratum radiatum in the CA1 area of the hippocampus.

To obtain an input/output (I/O) curve, short paired-pulse stimuli (50 ms interval, 5V pulse size, 0.1 ms pulse duration with bipolar electrode) with intensities ranging from 0.1 mV to 1.0 mV were delivered to evoke fEPSPs (Isolated pulse stimulator 2100, A-M systems, USA). The first response peak was used for plotting the I/O curve. Stimulus intensity corresponding to 45 – 50 % of the maximum response in the I/O curve was applied during the entire recording. fEPSPs were continuously recorded using a paired-pulse stimulation protocol given every 20 s with a 50 ms interstimulus interval.

Following 10 min of baseline recording, if the fEPSP responses are stabilized, a high-frequency stimulation (HFS, 100 Hz, 10 pulses repeated 4 times) protocol was applied to induce long-term potentiation. Signals obtained during recordings were amplified using a dual-channel extracellular amplifier (EXT-02F, NPI Electronic, Germany). To reduce noise, recorded signals were filtered with a 3-kHz low-pass filter and a 3-Hz high-pass filter. Signals were digitally amplified with a scale factor of 5 and a gain of 200 and stored till offline analysis (CED 1401, Cambridge electronic design, UK). Data were analyzed offline using Spike 2 (version 8.06, CED, United Kingdom) software. The mean value of 5 consecutive mean fEPSP slopes, each derived from three peaks obtained one step before, was computed and plotted on the graph. These mean values were used to represent fEPSP responses for every 5-minute interval, resulting in one data point on the graph per 5-minute period. At the end of recording, fEPSP responses were ablated using 10 μM tetrodotoxin (TTX).

After fEPSP recording, hippocampal slices were fixated in 2% glutaraldehyde and 1% formaldehyde in phosphate buffered saline for 24 hours, washed overnight in cacodylate buffer, postfixed with OsO4, dehydrated, and embedded in Epon (Roth, Germany). Semithin sections (1 um) were stained with toluidine blue. Ultrathin sections (50 nm) were stained with uranyl acetate and lead citrate and viewed with a Zeiss EM 900N electron microscope.

## Results

Among the 5 separate preparations of corticohippocampal slices, one preparation had apparent ice formation within slices already during cooling, and a second one had apparent recrystallization of ice within slices during rewarming. Both preparations did not yield any slice with recordable fEPSP response. The other three preparations yielded recordable fEPSP responses and are reported here. Untreated control slices of these successful preparations exhibited an appropriate stimulus intensity according to I/O-curve measurement of 0.2 mA. After vitrification, slices exhibited an appropriate stimulus intensity according to I/O-curve measurement of 0.5 mA and a reduced amplitude (approx. 0.4 mV). Response to HFS was recorded in one slice of the three preparations each, with two recordings showing HFS-induced potentiation of fEPSP slopes and normalization to baseline after 60 minutes, and one recording showing no response to HFS at all, albeit successful ablation with TTX after 60 minutes. fEPSP recording and responses are shown in Figure 1.

**Figure 1:**
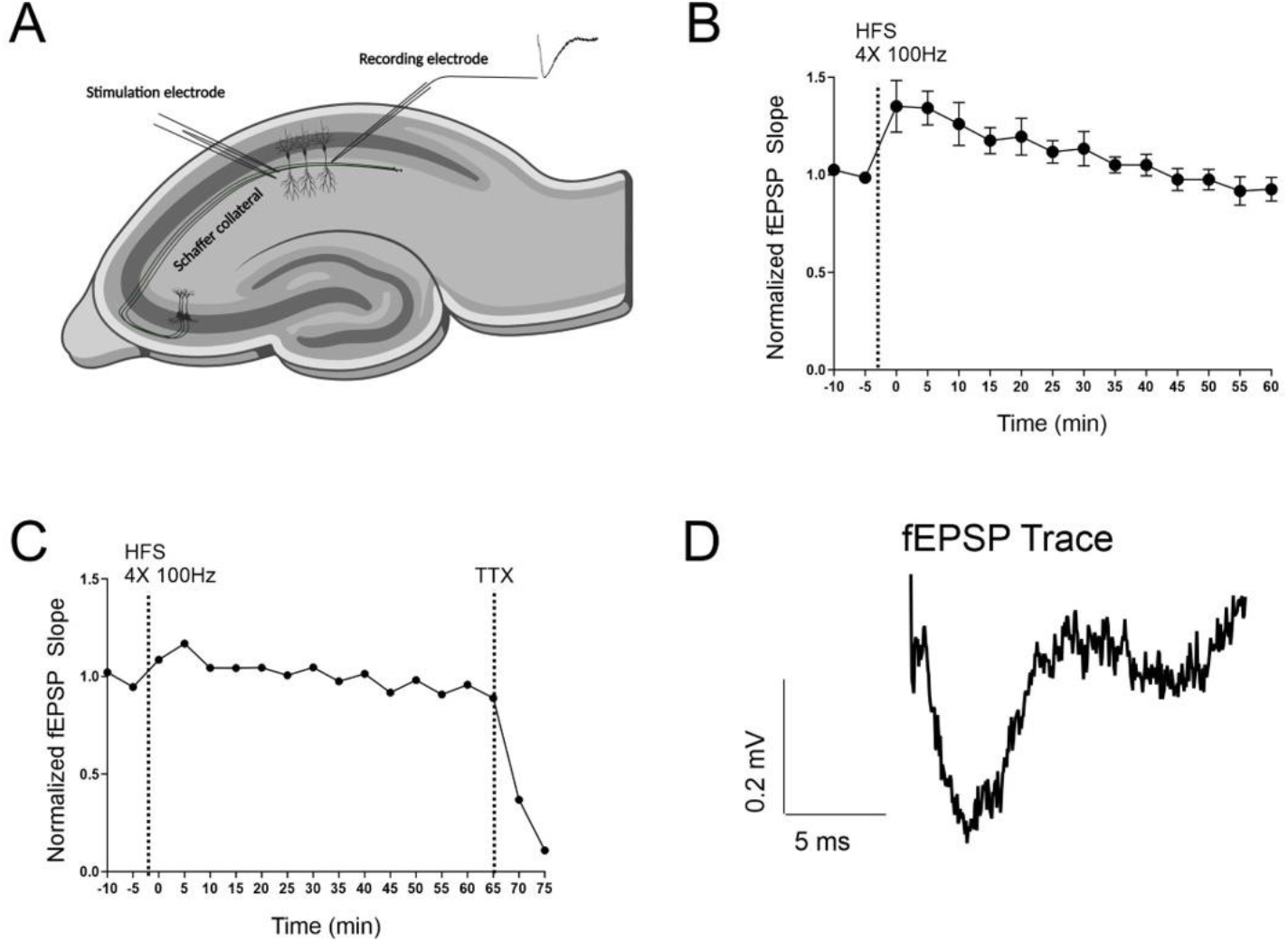
fEPSP recordings and responses. A: Stimulation of SC and fEPSP recoding in the stratum radiatum region in the CA1 area of the hippocampus. B: averaged fEPSP slopes for 10-minute baseline followed by HFS and recording for 60 minutes. C: fEPSP slopes lacking HFS response after 10 minutes and ablation with TTX after 65 minutes. D: representative fEPSP trace after vitrification. Toluidin blue semithin sections showed swelling in all layers of the hippocampus after vitrification, see Figure 2. Electron microscopy confirmed the appearance of cell swelling, and showed vacuolization and neuronal membrane ruptures with leakage of cytoplasm, see Figure 3.

**Figure 2.**
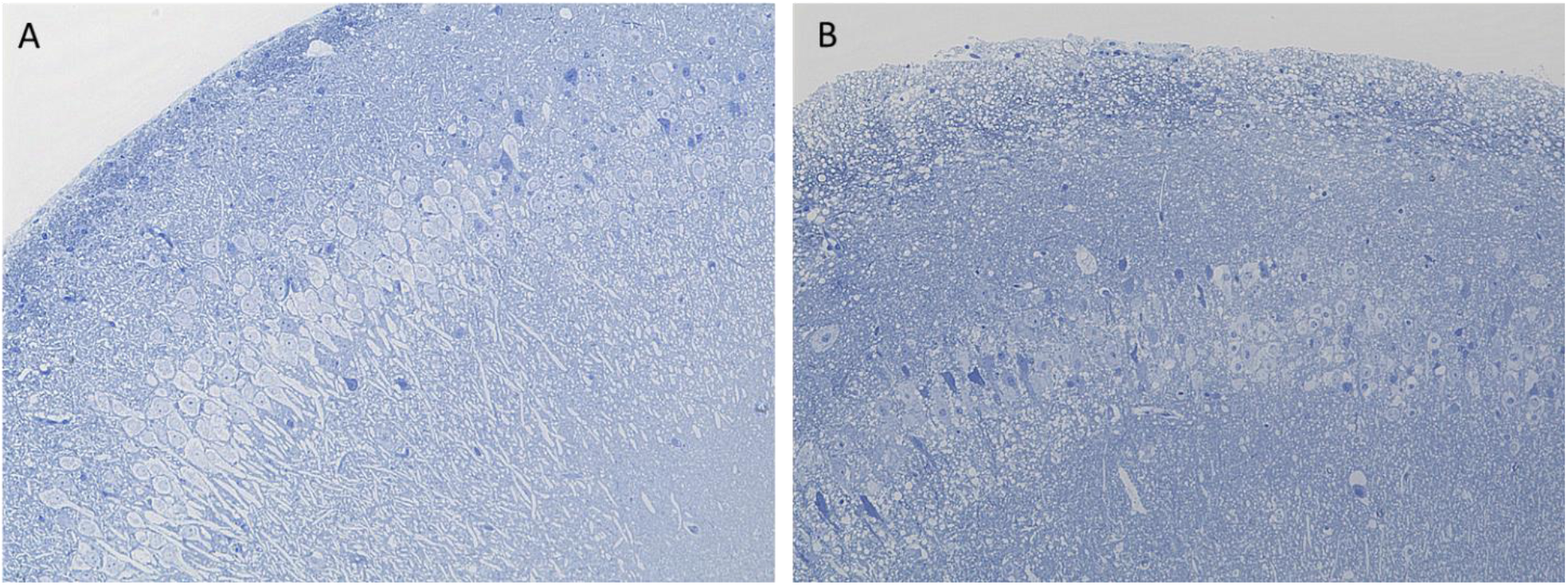
Light microscopy of Toluidin blue semithin sections of corticohippocampal slices after fEPSP recording. Cell swelling in the stratum oriens, pyramidale and radiatum of the hippocampus post-vitrification (B) compared to untreated control slices (A).

**Figure 3.**
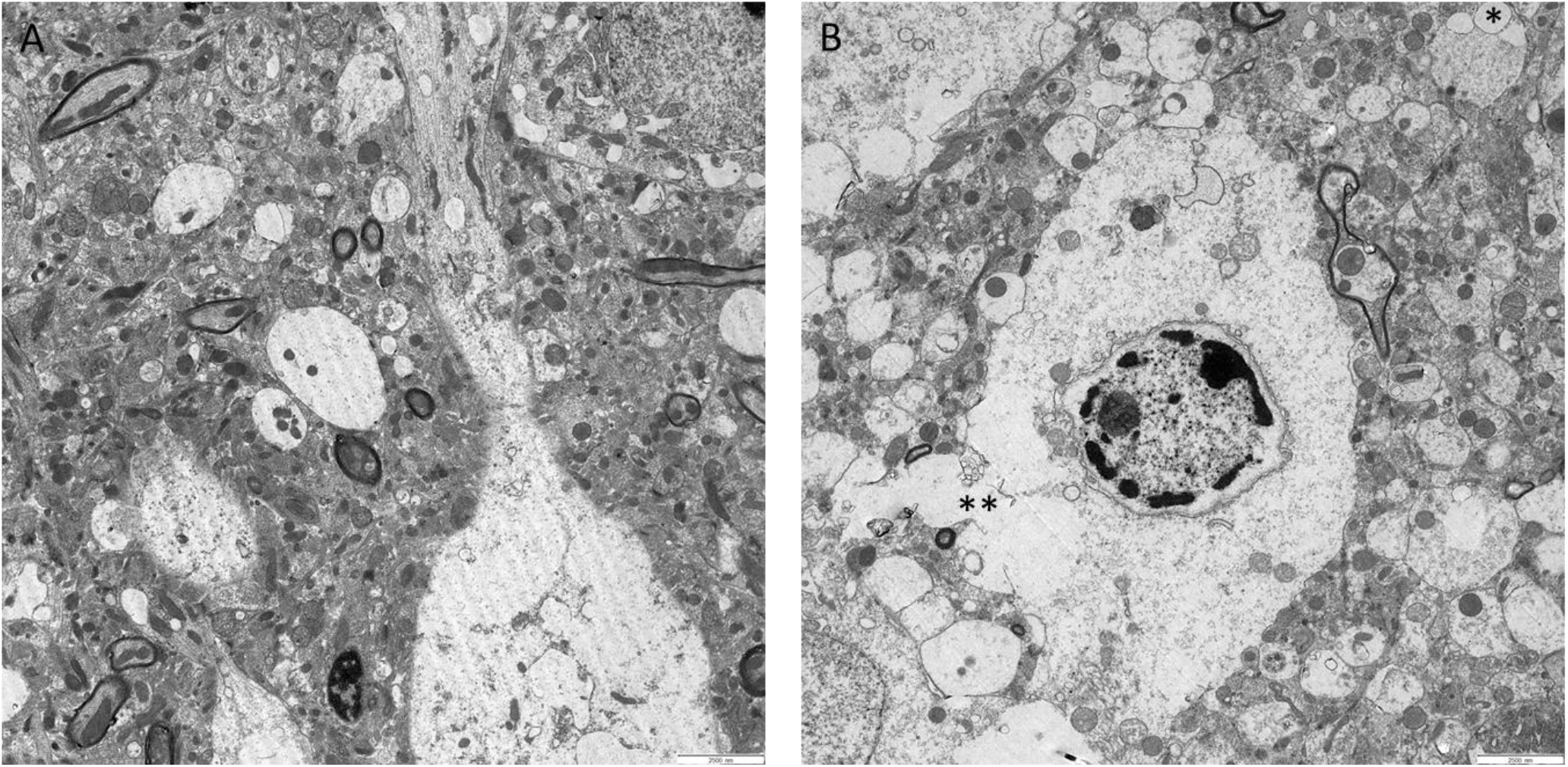
Electron microscopy of corticohippocampal slices in the stratum pyramidale of the CA2 subfield after fEPSP recording. Extensive swelling, vacuolization (asterisk) and cell membrane rupture (double asterisk) in the hippocampus post-vitrification (B) compared to untreated control slices (A). Magnification 4000x each.

## Discussion

Our results show that neural excitability and synaptic transmission can be recovered after vitrification. Successful potentiation of synaptic transmission by HFS also demonstrates retained synaptic plasticity after vitrification. Observation of HFS-induced potentiation and successful ablation with TTX suggests that the recorded fEPSP response is unlikely to be an artifact. However, fEPSP responses were pathological with reduced amplitude and increased stimulus intensity. This was reflected in structural alterations indicative of necrosis, i.e., swelling, vacuolization, and cell membrane rupture. The presence of necrosis in corticohippocampal slices with successful HFS-induced potentiation explains the failure of the induction of stable potentiation. This also highlights the need to combine functional with structural readouts. Recording of slices after cryoprotectant loading and unloading but not vitrification yielded similarly pathological fEPSP responses (data not shown). In the three cases of successful vitrification, cryoprotectant and hypothermia exposure thus appears to be the most important cause of toxicity for the corticohippocampal slices. Our technique is limited by low reproducibility with a success rate of only 60% of experiments. Reasons could be inadequately low cooling and warming rates, or mechanical damage in the vitrified state due to manual handling. In conclusion, we demonstrate that partly reversible deactivation of brain function by a cryopreservation technique is possible. Refining this technique could be of great interest to neuroscientific applications and beyond.

## Literature

1. Luyet, B.J. and M.P. Gehenio, Life and Death at Low Temperatures. 1940: Biodynamica.

2. Dubochet, J., et al., Electron microscopy of frozen water and aqueous solutions. Journal of Microscopy, 1982. 128(3): p. 219–237.

3. Dubochet, J., Cryo-EM—the first thirty years. Journal of Microscopy, 2012. 245(3): p. 221–224.

4. Armstrong, D.M., 13. The Nature of Mind, in Volume I Readings in Philosophy of Psychology, Volume I, B. Ned, Editor. 1980, Harvard University Press: Cambridge, MA and London, England. p. 191–199.

5. Golgi, C., Sulla struttura della sostanza grigia dell cervello’, Gazz. Med Lombarda, 1873. 33: p. 224–46.

6. Cajal, S.R.y., Revista trimestral de histología normal y patológica, Año 1, n. 1. Barcelona; Casa Provincial de la Caridad, 1888.

7. Brodmann, K., Vergleichende Lokalisationslehre der Grosshirnrinde in ihren Prinzipien dargestellt auf Grund des Zellenbaues. 1909: Barth.

8. Amunts, K., et al., BigBrain: An Ultrahigh-Resolution 3D Human Brain Model. Science, 2013. 340(6139): p. 1472–1475.

9. Motta, A., et al., Dense connectomic reconstruction in layer 4 of the somatosensory cortex. Science, 2019. 366(6469): p. eaay3134.

10. Loomba, S., et al., Connectomic comparison of mouse and human cortex. Science, 2022. 377(6602): p. eabo0924.

11. GoogleResearch, Google Research embarks on effort to map a mouse brain. 2023.

12. Lichtman, J.W. and J.R. Sanes, Ome sweet ome: what can the genome tell us about the connectome? Curr Opin Neurobiol, 2008. 18(3): p. 346–53.

13. Witvliet, D., et al., Connectomes across development reveal principles of brain maturation. Nature, 2021. 596(7871): p. 257–261.

14. Barranco, D., V. Cabo-Ruiz, and R. Risco, Use of fine capillaries for cryopreservation of Caenorhabditis elegans by vitrification. Cryobiology, 2023. 113: p. 104585.

15. Winding, M., et al., The connectome of an insect brain. Science, 2023. 379(6636): p. eadd9330.

16. Ohyama, T., et al., A multilevel multimodal circuit enhances action selection in Drosophila. Nature, 2015. 520(7549): p. 633–639.

17. Studer, D., B.M. Humbel, and M. Chiquet, Electron microscopy of high pressure frozen samples: bridging the gap between cellular ultrastructure and atomic resolution. Histochem Cell Biol, 2008. 130(5): p. 877–89.

18. Korogod, N., C.C.H. Petersen, and G.W. Knott, Ultrastructural analysis of adult mouse neocortex comparing aldehyde perfusion with cryo fixation. eLife, 2015. 4: p. e05793.

19. Lu, X., et al., Preserving extracellular space for high-quality optical and ultrastructural studies of whole mammalian brains. Cell Rep Methods, 2023. 3(7): p. 100520.

20. Luyet, B. and F. Gonzales, Growth of nerve tissue after freezing in liquid nitrogen. Biodynamica, 1953. 7(141-144): p. 171–4.

21. Robbins, R.J., et al., Cryopreservation of human brain tissue. Experimental Neurology, 1990. 107(3): p. 208–213.

22. Jensen, S., T. Sørensen, and J. Zimmer, Cryopreservation of fetal rat brain tissue later used for intracerebral transplantation. Cryobiology, 1987. 24(2): p. 120–134.

23. Otto, F., et al., Cryopreserved rat cortical cells develop functional neuronal networks on microelectrode arrays. Journal of Neuroscience Methods, 2003. 128(1): p. 173–181.

24. Heumüller-Klug, S., et al., Impact of cryopreservation on viability, gene expression and function of enteric nervous system derived neurospheres. Front Cell Dev Biol, 2023. 11: p. 1196472.

25. Ma, W., T. O’Shaughnessy, and E. Chang, Cryopreservation of adherent neuronal networks. Neuroscience Letters, 2006. 403(1): p. 84–89.

26. Uemura, M. and H. Ishiguro, Freezing behavior of adherent neuron-like cells and morphological change and viability of post-thaw cells. Cryobiology, 2015. 70(2): p. 122–135.

27. Suda, I., K. Kito, and C. Adachi, Viability of long term frozen cat brain in vitro. Nature, 1966. 212(5059): p. 268–70.

28. Suda, I., K. Kito, and C. Adachi, Bioelectric discharges of isolated cat brain after revival from years of frozen storage. Brain Research, 1974. 70(3): p. 527–531.

29. Pascoe, J.E. and A.S. Parkes, The survival of the rat’s superior cervical ganglion after cooling to –76° C. Proceedings of the Royal Society of London. Series B - Biological Sciences, 1957. 147(929): p. 510–519.

30. Pichugin, Y., G.M. Fahy, and R. Morin, Cryopreservation of rat hippocampal slices by vitrification. Cryobiology, 2006. 52(2): p. 228–40.

